# Nucleosome positioning sequence patterns as packing or regulatory

**DOI:** 10.1101/755272

**Authors:** Erinija Pranckeviciene, Sergey Hosid, Nathan Liang, Ilya Ioshikhes

## Abstract

Nucleosome positioning DNA sequence patterns (NPS) - usually distributions of particular dinucleotides or other sequence elements in nucleosomal DNA - at least partially determine chromatin structure and arrangements of nucleosomes that in turn affect gene expression. Statistically, NPS are defined as oscillations of the dinucleotide periodicity with about 10 base pairs (bp) which reflects the double helix period. We compared the nucleosomal DNA patterns in mouse, human and yeast organisms and observed few distinctive patterns that can be termed as packing and regulatory referring to distinctive modes of chromatin function. For the first time the NPS patterns in nucleus accumbens cells (NAC) in mouse brain were characterized and compared to the patterns in human CD4+ and apoptotic lymphocyte cells and well studied patterns in yeast. The NPS patterns in human CD4+ cells and mouse brain cells had very high positive correlation. However, there was no correlation between them and patterns in human apoptotic lymphocyte cells and yeast, but the latter two were highly correlated with each other. By their dinucleotide arrangements the analyzed NPS patterns classified into stable canonical WW/SS (W=A or T and S=C or G dinucleotide) and less stable RR/YY (R=A or G and Y =C or T dinucleotide) patterns and anti-patterns In the anti-patterns positioning of the dinucleotides is flipped compared to those in the regular patterns. Stable canonical WW/SS patterns and anti-patterns are ubiquitously observed in many organisms and they had high resemblance between yeast and human apoptotic cells. Less stable RR/YY patterns had higher positive correlation between mouse and normal human cells. Our analysis and evidence from scientific literature lead to idea that various distinct patterns in nucleosomal DNA can be related to the two roles of the chromatin: packing (WW/SS) and regulatory (RR/YY and “anti”).

**Author summary:** Precise positioning of nucleosomes on DNA sequence is essential for gene regulatory processes. Two main classes of nucleosome positioning sequence (NPS) patterns with a periodicity of 10bp for their sequence elements were previously described. In the 1st class AA,TT and other WW dinucleotides (W= A or T) tend to occur together in the major groove of DNA closest to the histone octamer, while SS dinucleotides (S= G or C) are primarily positioned in the major groove facing outward. In the 2nd class AA and TT are structurally separated (AA backbone near the histone octamer, and TT backbone further away), but grouped with other RR (R is purine A or G) and YY (Y is pyrimidine C or T) dinucleotides. In [8] we also described novel anti-NPS patterns, inverse to the conventional NPS patterns: WW runs inverse to SS, RR inverse to YY. We demonstrated that Yeast nucleosomes in promoters show higher correlation to the RR/YY pattern whereas novel anti-NPS patterns are viable for nucleosomes in the promoters of stress associated genes related to active chromatin remodeling. In the present study we attribute different functions to various NPS patterns: packing function to WW/SS and regulatory – to RR/YY and anti-NPS patterns.

## Introduction

Nucleosomes bring order to eukaryote genome and serve three primary functions [1]: provide measures of packaging and stabilize negative super coiling of DNA in vivo; provide epigenetic layer of information guiding interactions of trans-acting proteins with the genome through their histone modification; directly regulate access to the functional elements of the genome by their positioning. Nucleosomes have inhibitory effects on transcription by reducing access of transcription machinery to genomic DNA. Positioning and occupancy of nucleosomes contribute to the heterogeneity and flexibility of gene expression [2] and take part in chromatin activity. Biological consequences of nucleosome positioning and occupancy vary between cell types and conditions [3].

Negatively charged DNA backbone and positively charged histone octamer do not require DNA sequence specificity to make bonds. Core histone sequences are conserved among different species, therefore the biophysical principles of the histone assembly that determine histone preferences to certain DNA sequences should be universal across organisms [4]. However, it is thought that certain numbers of nucleosomes [5] in genomes are positioned by a preference of some DNA sequence patterns over the other. Certain features of the DNA sequence, such as in Widom 601 sequence, have much higher affinity to the histone octamer in vitro [6]. Such high affinity DNA sequence pattern is composed from WW (AA,TT,AT) dinucleotide steps in which a minor groove faces the histone octamer occurring at 10 base pairs (bp) periodicity. In addition, in between those steps where the major groove faces the octamer, the SS dinucleotides occur - in particular G or C followed by C or G [7], [8]. Within 147 bp that are wrapped around the histone octamer such periodical recurrence of distinctive dinucleotides facilitates a sharp bending of DNA around the nucleosome [10]. It is already known that specific compositions of dinucleotides make DNA more bendable [11] and that nucleosome linker regions show strong preference to sequences that resist DNA bending and disfavor nucleosome formation [12]. The GC rich nucleosome regions have higher nucleosome density while AT regions are more nucleosome depleted [13].

Generally, it is thought that nucleosome organization in encoded by a DNA sequence in eukaryotic genomes [14–17]. Specifically arranged nucleosome favoring and deterring sequences in vivo may act as containers attracting nucleosomes [1]. In vitro and in vivo nucleosome maps are noticeably different by the absence of periodically positioned and phased nucleosomal arrays in the in vivo maps where an ATP-dependent chromatin re-modeler is responsible for the well-spaced nucleosome array [18]. In addition to the sequence composition there are other factors in vivo such as post-translational modifications (PTM) of histone tails, transcription factor binding and remodeling complexes that influence positioning of nucleosomes and in turn tune chromatin accessibility and gene expression [5, 18].

It was shown that nucleosome remodeling takes place in response to stress and alters a gene expression. A single cell study performed on nucleosomes in yeast under glucose rich (PHO5 gene is silenced) and glucose starvation (PHO5 gene is expressed) conditions revealed that under starvation yeast cells lose nucleosomes in the PHO5 promoter [2]. However, mutants with enhanced AA/TT/TA periodicity in PHO5 gene promoter did not lose nucleosomes under starvation – the nucleosomes were positioned as in wild type under the nutrient rich conditions. This data showed that the periodicity enhancing mutations stabilized nucleosomes in such way that alienated chromatin re-modelers [2]. Another comparative study of nucleosomal DNA sequences in human CD4+ and apoptotic lymphocyte cells revealed a loss of GC rich nucleosomes in apoptotic cells [19]. Remodeling and reduced nucleosome occupancy was also observed in nucleus accumbens cells (NAC) of mouse brain in response to stress in chronic social defeat conditions [20]. Altered occupancy and positioning of nucleosomes was associated with a deactivation of genes implicated in stress susceptibility and also was significantly correlated with altered binding activity of TAP-utilizing chromatin assembly and remodeling factor-ATF complex.

Ensembles of nucleosomal DNA can differ between species and so the usage of nucleosomes in gene regulation [5]. Genomes are programmed to organize their own nucleosome occupancy. However, intrinsic histone preferences of specific k-mer sequences might be species specific [7, 21]. In higher eukaryotes genomic elements are closed by nucleosome unless active nucleosome displacement leads to the activation of this element. In unicellular organisms the genomic sites are open allowing transcription factor (TF) binding unless a nucleosome is actively repositioned there. Promoters of multicellular organisms are characterized by sequences favoring nucleosomes and in unicellular organisms by the disfavoring sequences [22].

Sequence based mechanisms governing nucleosome positioning and stability in different conditions and organisms genome-wide can be statistically characterized as patterns of periodically occurring k-mers in nucleosomal DNA [23]. The most prominent patterns observed in nucleosomal DNA across various organisms and particularly in yeast consist of the dinucleotides WW and SS (W=A or T and S=C or G) and RR and YY (R=A or G and Y=C or T) occurring at steps of average 10.1-10.4 base pairs that reflect the double helix period. A variety of such patterns was elucidated from nucleosomal DNA and shown that distinct pattern classes (termed pattern and anti-pattern) occur in nucleosomal DNA of nucleosomes modulating chromatin accessibility in promoters [8, 9, 24]. A signal of a periodical occurrence of specific dinucleotides in these patterns results from the averaging of a batch of nucleosomal DNA sequences and not necessarily this average pattern is found in any one given individual nucleosome. The analogy for such signal in a different biomedical domain is evoked potential of electroencephalogram which can be seen only from average signal of multiple experiments. The signals in nucleosomal DNA were shown not to be random by shuffling dinucleotides in nucleosomal DNA sequences [19].

We hypothesize that patterns originating from nucleosomal DNA in different experimental conditions may provide descriptive means to characterize and distinguish different states of chromatin function. In our study for the first time nucleosomal DNA patterns in nucleus accumbens cells (NAC) of stress-susceptible, stress-resilient and control mice brain [20] were characterized and compared to the dinucleotide frequency distribution patterns in human CD4+ and apoptotic lymphocyte cells. In these distinct organisms and very distinct biological conditions we aimed to discover fundamental similarities and differences in their nucleosomal DNA patterns and compared them to yeast. The nucleosomal DNA patterns in human and yeast cells were taken from the previously published studies [8, 19]. Comparing nucleosomal DNA patterns in mouse, human and yeast revealed that several distinct modes of chromatin function can be characterized by the few distinctive patterns which arguably can be named as packing and regulatory.

## Materials and methods

### Nucleosomal DNA patterns in human CD4+ and apoptotic lymphocyte cells

Sequences of nucleosomal DNA of +1 nucleosome in human CD4+ cells [25] in apoptotic lymphocyte cells [26] and their patterns characterized previously [19] were obtained from [19]. The original sequences of nucleosomes in human CD4+ cells (total 581507) and in apoptotic lymphocyte cells (total 711873) were flanked by 200bp on both sides of the dyad.

### Nucleosomal DNA patterns in nucleus accumbens cells of mouse brain

#### Original sequencing data

The original nucleosome sequencing data of mouse brain NAC cells [20] were obtained from NCBI GEO archive under accession GSE54263. The short read sequence files accessioned by SRR113826[1-9] were downloaded from Short Read Sequences Archive (SRA) by NCBI SRA toolkit as fastq files. The fastq files contain paired MNase-seq (MNase digested histone H3) reads sequenced on Illumina HiSeq with 99bp read length in three biological replicates of control, stress-resilient and stress-susceptible mice from [20].

#### Alignment

The reads were aligned to the mouse reference genome *mm9* by bwa mem [27]. The 95%-98% reads were uniquely aligned and overall genome-wide mean coverage was 4 reads. The bam files were converted into bed format and genome-wide coverage was computed by BEDTools [28]. Mean length of covered regions was 279.29 ± 8.45 and mean length of zero-coverage regions was 99.48 ± 12.52.

#### Determination of nucleosome sequences

Sequences of nucleosomal DNA were obtained using algorithm as in [19] from information in the genome-wide coverage profile. The coverage profile peaks identify start positions of nucleosomes [29]. Genomic positions in which peaks attain maximum were identified by applying Gaussian smoothing to the coverage profile and taking a position in which smoothed profile has maximum. The choice of smoothing window depends on the data. We investigated several window sizes. Optimal size for this data was 70bp which is also a recommended window size for this type of data [30]. Peaks that are twice of average coverage [29] usually identify nucleosome positions. In this data average coverage was 4, therefore genome-wide peak summits attaining a height less than 10 were discarded. Genomic coordinates of nucleosome bound sequences were calculated from the mapped reads overlapping the summit positions such that a distance between the summit and either end of the read is no less than 30 bp. As in [31] The 5’ start positions of the summit overlapping reads were extended by 20 bp upstream and the 3’ end positions by 100 bp downstream flanking the146 bp nucleosome and it dyad position from both sides. The upstream flanking is shorter since it is expected to capture a cleavage site which will determine a most likely nucleosome start position. The DNA sequences of these intervals were extracted from mouse mm9 reference genome by BEDTools, aligned by the experimental 5’-end and formatted into fasta files. The nucleosome sequences of the first best phased nucleosome within 1000bp downstream of gene TSS were retained for further analysis. The mm9 gene TSS coordinates were downloaded from UCSC Genome Table Browser [32]. Variable number of sequences and associated RefSeq genes were obtained in three replicates in each biological condition. On average from 6 to 8 sequences aligned by the experimental 5’ end represented a well phased nucleosome in the downstream vicinity of RefSeq gene TSS. Table 1 provides summary of the aligned raw data and sequences. To analyze patterns in the nucleosomal DNA in control, susceptible to stress and resilient to stress mice the sequences from the biological replicates in each condition were aggregated into one fasta file.

**Table 1.** Summary of aligned GSE54263 original sequences from [20] and total counts of RefSeq genes and sequences of first well phased nucleosome within 1000bp downstream of mm9 Refseq gene TSS.

| SRA name | Condition | GEO ID | Raw reads | Uniquely aligned | Nucleosome sequences | RefSeq Genes |
| --- | --- | --- | --- | --- | --- | --- |
| SRR1138261 | control | GSM1311267 Con_H3 | 112,379,273 | 110,283,095 | 58658 | 8418 |
| SRR1138262 | control | GSM1311267 Con_H3 | 127,059,884 | 122,696,395 | 65909 | 9294 |
| SRR1138263 | control | GSM1311267 Con_H3 | 120,053,422 | 116,102,876 | 85433 | 11860 |
| SRR1138264 | resilient | GSM1311268 Res_H3 | 102,811,190 | 99,612,874 | 40464 | 5775 |
| SRR1138265 | resilient | GSM1311268 Res_H3 | 124,676,340 | 120,399,602 | 65642 | 9084 |
| SRR1138266 | resilient | GSM1311268 Res_H3 | 127,632,626 | 123,374,019 | 93327 | 12865 |
| SRR1138267 | susceptible | GSM1311269 Sus_H3 | 105,995,184 | 102,501,729 | 42631 | 5985 |
| SRR1138268 | susceptible | GSM1311269 Sus_H3 | 115,654,263 | 111,596,579 | 57352 | 7940 |
| SRR1138269 | susceptible | GSM1311269 Sus_H3 | 130,984,076 | 126,572,005 | 98886 | 13661 |

#### Computation of patterns in nucleosomal DNA sequences

Patterns of positional dinucleotide frequency distributions along nucleosomal DNA sequences from a bulk of sequences were computed utilizing a previously described algorithm [19]. Given a binary matrix of dinucleotide occurrences in sequences coded as 1 and else as 0 a pattern of dinucleotide frequency of occurrence is a sum of occurrences of the selected dinucleotide at every position along a sequence normalized by a number of sequences. Peaks in dinucleotide frequency patterns along the nucleosomal DNA sequence have a recognizable dyad-symmetry [33]. The dyad-symmetry feature helps to determine a nucleosome’s position in a bulk of sequences. At the nucleosome position centered on dyad the dinucleotide distribution patterns on forward and reverse complementary strands will have a maximum positive correlation. To determine nucleosome’s position Pearson correlation between the dinucleotide frequency distribution profiles on forward and reverse complementary strand is computed at each position of the nucleosome sequence within a sliding 146 base pair long window. Positions in which Pearson CC attains maximum for each dinucleotide are examined. Identification of the nucleosome position can’t be fully automated and has to be verified for proximity to a cleavage site, because of varying nature of correlations across conditions. Once the nucleosome’s position is identified the corresponding computed dinucleotide patterns are symmetrized. Subsequently composite patterns of Weak-Weak/Strong-Strong WW/SS (W=A or T, S=C or G) and Purine-Purine/Pyrimidine-Pyrimidine RR/YY (R=A or G, Y= C or T) dinucleotides are computed. The patterns can be normalized and smoothed to reduce small noisy peaks to compute their periodograms. This algorithm is implemented as dnpatterntools v1.0 suite of binary C++ programs and shell scripts. By using dnpatterntools we reproduced previously determined patterns in human cells and computed new patterns in mouse. More details on methods used in this study are outlined in S1 Appendix.

#### Patterns in yeast

The patterns in yeast’s nucleosomal DNA used in this study were obtained from [8].

#### Availability of tools and data

The dnpatterntools v1.0 software and example fasta files of processed mouse sequences are available from GitHub (https://github.com/erinijapranckeviciene/dnpatterntools). An example of analysis workflow in Galaxy [34] using a toy control mouse nucleosomal DNA as an example is freely available from dockerized dnpatterntools-galaxy instance from the docker hub (https://hub.docker.com) in a repository dnpatterntools-galaxy based on a galaxy-stable base image (https://zenodo.org/record/2579276).

## Results

### Baseline and periodicity

DNA sequences in nucleosomes are characterized by a periodical 10 base pairs AA/TT dinucleotide frequency distributions most clearly manifesting in yeast. At various extents it is seen also in other model organisms – human, mouse, worm [35] and fruit fly [36]. We investigated patterns formed by WW/SS and RR/YY pairs of dinucleotides and individual AA/TT/TA and CC/GG dinucleotides which carry nucleosome positioning signals by their periodical arrangements. Frequency analysis (Fourier transform) of dinucleotide profiles in mouse brain NAC showed peaks at 10.1-10.4 base period in agreement with the anticipated 10bp periodicity. Dinucleotide frequency distributions in mouse and human cells had a slightly different baseline possibly because of G+C content differences in human (41%) versus mouse (42%) genome. We observed very different pattern of nucleosome sequence composition in apoptotic cells compared to the pattern in mouse and normal human cells, suggesting a different sequence-based mechanism governing dynamics of nucleosomes in cells undergoing apoptosis. How the patterns compare in mouse brain NAC, human CD4+ and apoptotic cells is shown in Section 1 Figures 1-5 in S2 Appendix.

**Fig 1.**
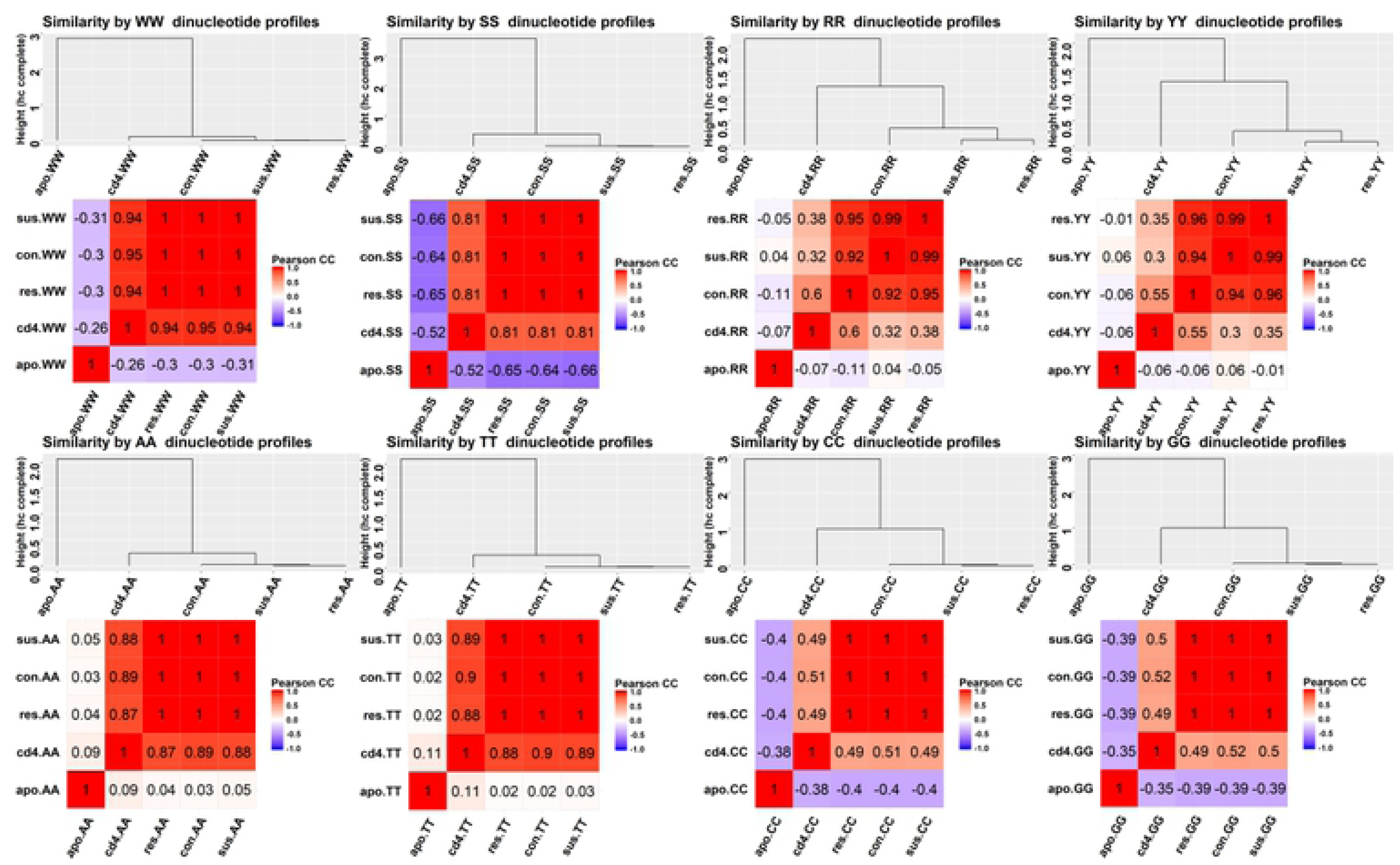
Similarities between dinucleotide patterns in nucleosomal DNA in mouse and human cells. For each composite (WW, SS, RR, YY) and individual (AA, TT, CC, GG) dinucleotide a similarity between the patterns in human CD4+ cells as cd4, human apoptotic lymphocyte cells as apo and NAC cells of mouse brains in control as con, stress-resilient as res and stress-susceptible as sus is shown. The similarity is quantized by a Pearson correlation coefficient a magnitude of which is shown in each cell of correlation matrix. Two representations of similarity are depicted for each dinucleotide: a heatmap in which color-coded cells show a Pearson correlation coefficient between two profles and color provides a direction - red is positive, blue is negative. The similarity dendrograms are computed from the correlation matrix shown in heat map and illustrate hierarchical complete linkage clustering of the patterns.

### Comparison of nucleosomal DNA patterns between human and mouse

Global similarity and difference between the dinucleotide frequency profiles in different conditions was quantified by computing Pearson correlation coefficient between the patterns pair-wise. Fig 1 shows heatmaps of correlation magnitudes between nucleosomal DNA patterns of human and mouse and dendrograms derived from these correlation matrices.

The patterns in normal human CD4+ cells and mouse brain NAC are strikingly similar even though their original experimental and functional contexts are very different. On the contrary, the patterns in apoptotic human cells were very different from those in mouse NAC and human CD4+ cells. However a high degree of similarity exists between the apoptotic patterns and the patterns observed in yeast [8]. The WW, AA and TT patterns in mouse NAC and human CD4+ cells have significant positive correlations (> 0.8). The CC, GG and SS patterns are less correlated (> 0.49). In all cases, the dinucleotide patterns in apoptotic cells have either no correlation or a negative correlation (WW is < –0.27 and SS is < –0.5) with the patterns in mouse and human CD4+ cells.

### Peak synchronicity and positional preferences

Positional preferences of nucleosomes to some sequence signatures manifest statistically as peaks of certain dinucleotides at certain positions in the frequency profiles computed from a large set of aligned sequences of nucleosomal DNA. We observed the maxima and minima (peaks and valleys) of AA/TT, CC/GG, WW/SS, and RR/YY dinucleotide patterns in mouse and human cells occur at a very high proximity. Positions in which a majority of dinucleotides in sequences from different conditions have well defined peaks can indicate *hubs* that may have special meaning in nucleosome dynamics. To locate hub positions, we use dinucleotide profiles transformed into a unit indicator sequence in which a +1 specifies a position of a maximum and a -1 specifies a position of a minimum and all other positions have zero values. We observed that the peaks mostly co-occur at intervals separated by a step of 10 ±1 base pairs. The sites in nucleosomal DNA in mouse and human cells were used differently by dinucleotides. In human cells the positions of multiple coinciding peaks along the nucleosomal DNA were ±69, ±47, ±46, ±43, ±28, ±23, ±19, ±13, ±7. In mouse cells they were: ±64, ±60, ±44, ±39, ±16, ±11. The human and mouse organisms also differed by identities of simultaneously occurring dinucleotide peaks.

Mutually exclusive dinucleotides such as WW and SS can’t simultaneously occupy same position. However, we observed that the peaks of CC and GG and the peaks of TA and YY simultaneously occur in mouse. In human CD4+ cells simultaneously occurring peaks were TA and GG and WW and SS. This means that analyzed sequences of nucleosomal DNA contain several different pattern classes. The single cell analysis study [2] showed that at any single moment single cells are in binary status: some have nucleosomes remodeled and some have not. This may explain presence of incompatible dinucleotide peaks in aligned nucleosomal DNA sequences from the same source. In human CD4+ cells nucleosomal DNA sequences formed two distinct clusters each favoring either WW or SS patterns as shown previously [19]. The *hub* sites in which a majority of dinucleotides including incompatible had peaks were ±45, ±31, ±20. These *hub* sites are located in ±2, ±3 and ±4 super helical locations (SHL) at which DNA interacts with histone octamer in a processes of remodeling that either stabilizes or destabilizes nucleosomes [37]. More details on synchronization of peaks, dinucleotides and the *hub* positions are provided in Section 2, Figures 6,7,8 and Table 1 in S2 Appendix.

### Structural signatures of dinucleotide patterns in various conditions

We investigated further how nucleosomal DNA patterns in higher human and mouse organisms compare to the well-established patterns [8] in unicellular yeast organism. As an additional dimension to our comparison we added a generalized distribution of superhelical locations (SHL). The SHL comprising minor and major grooves in nucleosomal DNA were derived by Cui and Zhurkin from roll angles of crystal structures of nucleosome core particle (NCP) illustrated in Fig. 3 in [11]. We overlapped the distributions of WW/SS, RR/YY, AA/TT/TA and CC/GG peaks in three organisms (yeast, mouse and human) on a map of SHL zones from the cartoon in Fig. 3 in [11] that corresponds to a half of nucleosomal DNA. A colour-coded ladder diagram of WW/SS and RR/YY peaks is shown in Fig. 2. The WW and SS steps and the RR and YY steps are represented opposite to each other to make it it easier to see an order in arrangement of these steps.

**Fig 2.**
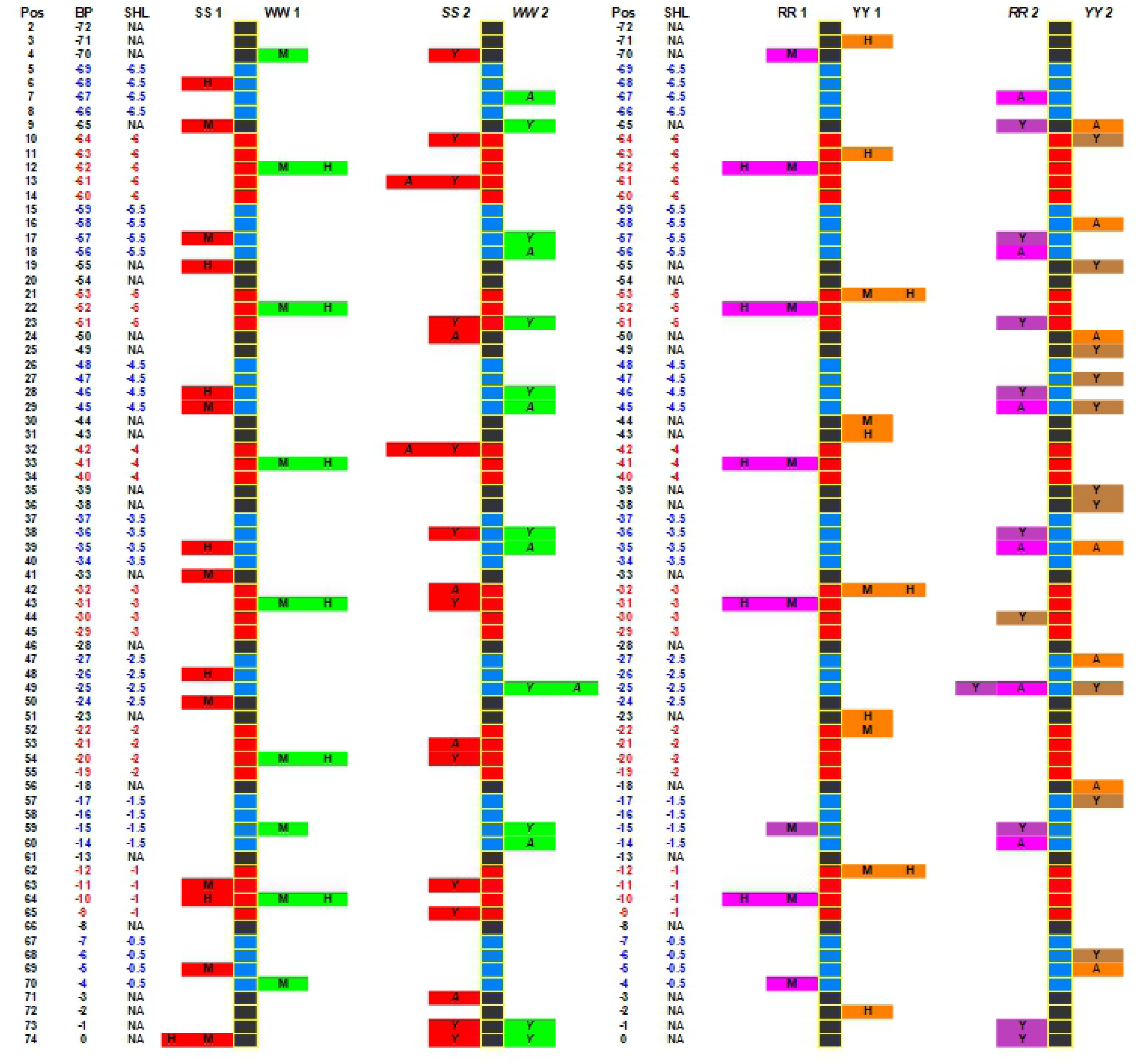
Distribution of WW/SS and RR/YY peaks in human, mouse and yeast. The ladder diagrams represent half of nucleosome with a dyad position at the bottom and proceeding up from -1 base pair position to the -72 base pair. The positions corresponding to the major grooves are shown in red and the minor are blue (as in cartoon of Fig.3 in [11] labeled by SHL numbers. The WW peaks are coded in green, the SS in red, the RR are coded in magenta shades and YY in orange shades. Each indicated peak inside the cell is labeled by a letter indicating which organism it belongs to (H- for human CD4+ cells, M- for NAC cell of control mouse, Y yeast and A for human apoptotic lymphocyte cells). The SS1/WW1 and RR1/YY1 diagrams show the peak distributions in human and mouse. The SS2/WW2 and RR2/YY2 diagram show peak distributions in yeast and human apoptotic cells. The numbers 1 and 2 designate these separate pattern classes. The ladder diagrams illustrate a synchronization between the peak occurrences. The WW peaks in human and mouse have SS peak counterparts in yeast and apoptotic cells and vice versa. The RR1/YY1 patterns appear to be very regular. The RR2/YY2 has less regular appearance in which peaks are mostly concentrated in SHL zones ±2.5, ±3.5 and ±4.5.

**Fig 3.**
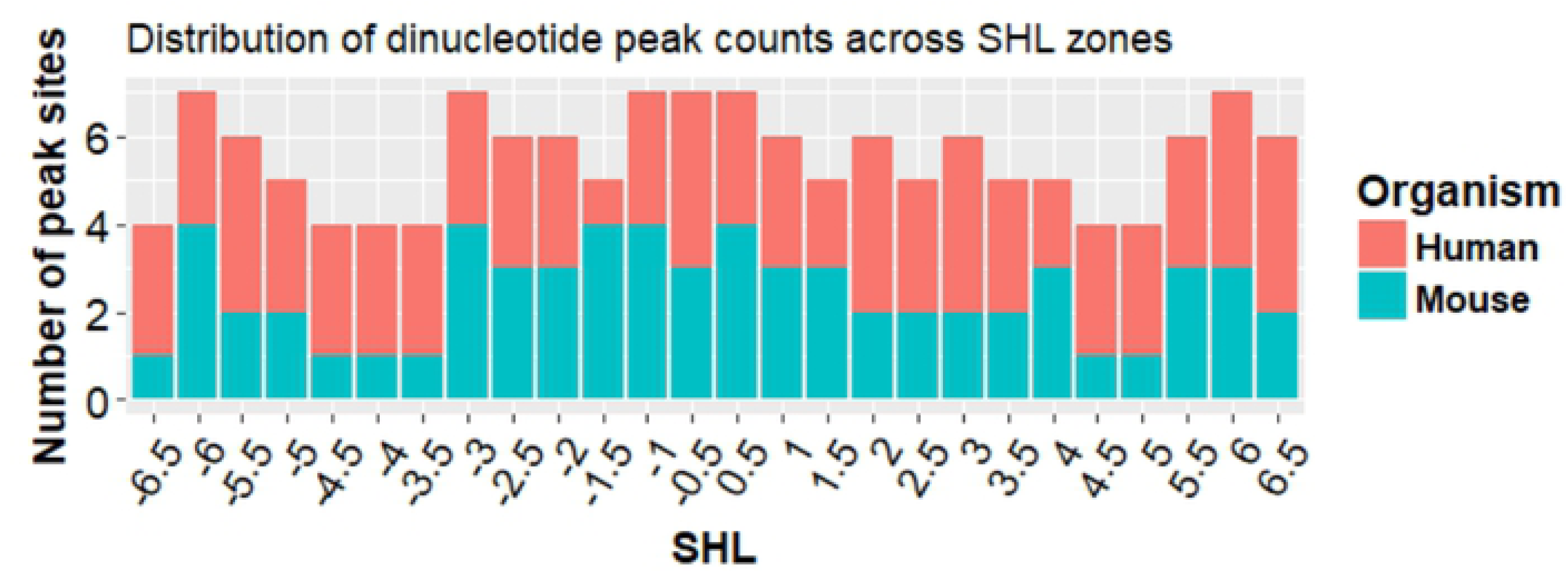
Comparison of peak counts across SHL zones in human and mouse. The bars show number of counts of peak sites across SHL in human CD4+ and apoptotic lymphocyte cells compared to the counts in mouse NAC in control, susceptible and resilient to stress mice.

In human CD4+ cells and mouse NAC cells most of WW peak sites coincide and are located in the major grooves. However, the WW peaks in yeast and human apoptotic cells mostly appear as opposite with respect to the same peak in human CD4+ cells and mouse NAC. The WW peaks in human CD4+ cells and mouse NAC occur in SHL major groove zones ±6, ±5, ±4, ±3, ±2, but in yeast and human apoptotic cells have SS peaks in these zones. And the opposite: the zones ±4.5 and ±2.5 have SS peaks in human CD4+ cells and mouse NAC, but in yeast and human apoptotic cells these locations contain WW peaks. The SS peaks proximal to the dyad in ±1.5 occur in mouse only. Most SS peaks in mouse and human are apart by ±1, ±2 base pairs but the central SS peak at the dyad coincides in both organisms. Generally, in all organisms peaks in SHL zone -1 have a smaller magnitude. Mouse and human at the dyad have SS peaks and yeast has both: SS and WW peaks. There are significant similarities of positional peak distributions in mouse NAC and normal human CD4+ cells. However, positional peak distribution in human apoptotic is significantly different and highly resembles peak distribution in yeast.

The distributions of RR/YY peaks have similar characteristics in human CD4+ and mouse. However, it is different in yeast cells and considerably different in human apoptotic cells. The RR2/YY2 pattern in yeast has RR (Purine-Purine) peaks occurring in SHL minor zones interchanging with YY (Pyrimidine-Pyrimidine) steps occurring in all SHL zones; whereas in human CD4+ cells and mouse NAC both the RR and YY steps occurr in major zones and are arranged in close proximity opposite to each other. The RR2/YY2 pattern in human apoptotic cells has a regular structure which is very different from other patterns. The observed WW/SS pattern configuration is a classical nucleosome favoring sequence configuration [7, 8]. As to observed RR/YY patterns – DNA structures containing these configurations of dinucleotides do not persist because of stiffness of RR and YY dinucleotide steps [8] and [39]. Peak site localization in SHL zones differ between mouse (control, susceptible and resilient to stress mice) and human (CD4+ and apoptotic cells). All sites that had either one or multiple peaks were counted in each SHL zone for mouse and for human. The distribution of counts is shown in Fig. 3. The peak positions of all dinucleotides in this study and their corresponding structural zones are summarized in Table 1 in S1 Tables.

Peak counts across SHL in human and mouse had a statistically significant difference (paired Wilcoxon rank sum test p-value =0.02). The SHL zone -1.5 have 4 peak sites in mouse but in human it has just one peak site in the apoptotic cells. The SHL zones ±4 ±4.5 and -3.5 in human have more peak sites compared to mouse. In SHL -6.5 zone only susceptible to stress mouse had a peak. SHL zones ± 1.5, ±4 and ±4.5 that differ between human and mouse in terms of positional preferences play roles in nucleosome dynamics. Most of DNA deformations take place in the SHL ± 1.5 zone. It has an increased binding of DNA proteins and it is most susceptible to DNA damage [38]. Other important zones in which multiple peaks from all organisms were observed are SHL zones ±2, ± 3 and ± 4. The SHL ± 2, ±3 are zones in which chromatin remodelers interact with DNA and in SHL ±4 zone histone H2A tail interacts with DNA which makes it a less stable zone [37]. Statistically defined relations between nucleosome positioning and covalent chromatin modification were noted – but there is no relationship established between histone modifications and specific nucleotides observed at a given location [5]. It is not clear yet how specific composition of DNA sequence might interact with these processes and affect nucleosome positioning and remodeling. A brief summary of current knowledge about importance of specific SHL zones trying to provide more context to the regularities observed in this work is provided in Section 3 in S2 Appendix.

## Discussion

We performed extensive descriptive and comparative study on dinucleotide patterns in nucleosomal DNA originating from cells in very different conditions: human CD4+ cells and human apoptotic lymphocytes, and NAC from brains of susceptible to stress, resilient to stress and control mice. The found patterns were compared to patterns in yeast. We used already published patterns for yeast and human organisms and characterized patterns in mouse NAC for the first time. Essentially, we aimed to find how patterns of nucleosome sequences are different or similar in these organisms and found important regularities.

Statistically, dinucleotide patterns in nucleosomal DNA provide information about sequence preferences of histones in forming nucleosomes and packing DNA. Patterns in this study were computed from a multiple sequences of the best phased most proximal to gene TSS nucleosomes genome-wide. Since the best phased closest to TSS nucleosome (+1 nucleosome) is one of primary factors determining how the rest of the nucleosomes will assemble, we hypothesized that patterns characterizing the nucleosomal DNA of these nucleosomes genome-wide in different organisms and conditions may reflect sequence patterns affecting chromatin function and packaging.

In response to stress nucleosomes undergo remodeling to alter expression of response genes. The remodeling may comprise various events: nucleosomes may change their position or occupancy, they may be evicted or otherwise change their configuration. The patterns derived from sequences of nucleosomes sequenced in a specific experimental condition (i.e. after remodeling) will represent state of chromatin that is specific to that condition because these sequences to some extent statistically represent the general pattern in sequence that attracted histones and formed nucleosomes. However, as it was shown, a remodeling may occur in a fraction cells where in a remaining fraction the original nucleosome configuration hasn’t changed. Therefore, in nucleosomal DNA sequence patterns there will be a mixture of patterns likely corresponding to the changed and unchanged condition and it is challenging to elucidate a very clear patterns. Nevertheless, some general trends can been observed.

We discovered that patterns in mouse nucleosomal DNA in control, stress-susceptible and stress-resilient conditions are almost identical. The stress affected mice differ slightly from a control in a spatial distribution of peaks in RR/YY patterns. The WW peaks in nucleosomal DNA in normal human CD4+ cells and in mouse NAC coincide and these peaks are located in a major grove SHL zones while the SS peaks are located in the minor grove SHL zones. We also observed that the patterns of WW and SS peak arrangement in human apoptotic cells are very similar to the patterns in yeast. The peaks in human apoptotic cells and in yeast are arranged in opposite way to the peaks observed in human CD4+ cells and mouse NAC. Namely, the WW maxima in human and mouse correspond to the SS maxima of yeast and human apoptotic cells and vice versa - the SS maxima in human CD4+ and mouse correspond to the WW maxima in yeast and human apoptotic cells. Namely, these patterns are inverse of each other. It is known that the WW/SS pattern as in yeast is the one that is favored by nucleosome formation: it is represented by stretches of WW dinucleotides (AA/TT/TA) on the face of the helical repeat that can directly interact with the histone. If this DNA sequence is altered to fit 5-bp intervals of AA/TT/TA dinucleotides with GC dinucleotides, then the nucleosome occupancy has been observed to increase dramatically [7]. This pattern – WW with SS inserted each 5 base pairs is characterized as a very stable. It facilitates a rotational positioning and has been observed in nucleosomal DNA from chickens, yeast, fruit flies, nematodes and humans, suggesting that the structural rules for rotational positioning are the same across species [11].

In this study we observed that WW dinucleotides in yeast and human apoptotic cells are located in minor grove SHL zones, however human CD4+ cells and mouse NAC in those zones have SS dinucleotides. It was stated [39] that CC and GG dinucleotides have more influence to nucleosome formation. Our investigated organisms may have employed different nucleotides, however a spatial structure of the dinucleotide arrangement is still preserved. The existence of the patterns in which WW/SS dinucleotides are used in opposite ways termed pattern and anti-pattern was already predicted [8] and investigated in promoters of mammalian cells [9]. Here the pattern corresponds to a spatial distribution of WW/SS peaks as in yeast and human apoptotic cells (represented by WW2/SS2 ladder in Fig. 2) and the anti-pattern corresponds to that in human and mouse (WW1/SS1 in 2). It is also known that chromatin becomes highly condensed during apoptosis [40]. Since these WW/SS patterns are very stable and they are universally used in many species and also seen in human apoptotic cells they could be termed as packing.

The RR and YY in human, mouse and yeast alternate in 3 to 5 base pair steps and again the RR peaks in human and mouse patterns occur in major grove SHL zones, while in yeast they are in minor groove SHL zones. The RR/YY patterns in this study are in agreement with (i) the already characterized RR/YY patterns in which dinucleotides are by 5 bases apart and (ii) in arrangement of peaks agree with the universal linear nucleosome positioning pattern YRRRRRYYYYYR [41]. The RR and YY dimers appear to be the most rigid dinucleotides, and therefore a DNA fragment consisting of the interchanging oligo R and oligo Y blocks that are 5–6 base pairs long should manifest a spectacular curvature in solution [42]. The RR/YY pattern in human apoptotic cells is very regular in which the RR and YY peaks occur in a close proximity to each other carrying a high resemblance to the in vitro Group 1 pattern of YR/YYRR motifs at sites SHL ±3.5 and ±5.5 described by Cui and Zhurkin [11]. This pattern facilitates severe DNA deformations at those sites and the positioning of nucleosomes is likely to be determined by interactions between H2A/H2B and DNA at those sites. Because of the high DNA flexibility imposed by the RR/YY patterns and because of the important functional consequences of SHL zones associated with these patterns these RR/YY patterns could be termed as regulatory.

## Conclusions

One of the novel contributions of this study to the nucleosome field is that we characterized patterns in mouse NAC for the first time. In addition we contributed software with which already characterized and published patterns can be reproduced and new patterns computed given fasta sequences of nucleosomal DNA. We described regularities observed in distributions of peaks in patterns on nucleosomal DNA in human, mouse and yeast and showed how these regularities in the arrangements of peaks classify dinucleotide patterns into WW/SS and RR/YY patterns and anti-patterns reported previously. We observed that multiple coinciding peaks in different patterns observed in different organisms and conditions are located in the nucleosome’s SHL zones that have important roles in chromatin functioning.

Striking similarities were found between the WW/SS patterns in yeast and human cells under the conditions of apoptosis. Biological significance of this finding is unknown. However, taking into account that WW/SS is a very stable pattern that is strongly favoring formation of nucleosomes and also that chromatin under apoptosis becomes highly condensed, we argue that it may suggest existence of several modes of chromatin functions that can be characterized by a few classes of sequence patterns that may be related to the two chromatin roles: packing and regulatory.

Based on our results and evidence from scientific literature we draw following conclusions:

- Yeast bulk nucleosome sequences display and tend to be mapped by canonical stable WW/SS patterns.
- Nucleosome sequences in mouse and human display and tend to be mapped by the canonical WW/SS anti-pattern and the RR/YY pattern.
- Promoter sequences in yeast tend to be mapped by the canonical RR/YY pattern and promoters of the yeast stress-related genes tend to be mapped by the canonical RR/YY anti-pattern [8].
- Patterns in nucleosomal DNA are related to the two roles of the chromatin: packing (WW/SS) and regulatory (RR/YY and “anti”).
- Our hypothesis for the future studies is that packing patterns tend to be preferred by evolutionary lower organisms and regulatory – by higher organisms.

## Supporting information

**S1 Appendix. Additional methodological details and examples of workflows to obtain nucleosome sequences and patterns.**

**S2 Appendix. Additional details on nucleosomal DNA patterns and superhelical locations.** This Appendix presents multiple plots of patterns in nucleosomal DNA of mouse and human with additional analysis and details on distributions of peaks. It contains a short summary on importance of superhelical locations.

**S1 Tables. Dinucleotide patterns and peak distributions.** Tables in this spreadsheet comprise:

- Table 1. Distributions of dinucleotide peaks in human mouse and yeast.
- Table 2. Dinucleotide patterns in human and mouse.
- Table 3. Comparison of peaks between human and mouse. Complementary to Fig. 3.
- Table 4. Excel representation of peak distributions in human, mouse and yeast patterns as a ladder. Complementary to Fig. 2.

### Acknowledgments

The University of Ottawa Faculty of Medicine Bridge Fund 2015-2016 to I.I. is greatly acknowledged. The authors thank Prof. Eric Nestler for providing NAC data prior to publication.

